# Oligodendrocytes in human iPS cell-derived cortical grafts remyelinate adult rat and human cortical neurons

**DOI:** 10.1101/2023.03.24.534133

**Authors:** Raquel Martinez-Curiel, Linda Jansson, Oleg Tsupykov, Natalia Avaliani, Constanza Aretio-Medina, Isabel Hidalgo, Emanuela Monni, Johan Bengzon, Galyna Skibo, Olle Lindvall, Zaal Kokaia, Sara Palma-Tortosa

**Affiliations:** Laboratory of Stem Cells and Restorative Neurology, Lund Stem Cell Center, Lund University, SE-22184, Lund, Sweden; Department of Cytology, Bogomoletz Institute of Physiology; Institute of Genetic and Regenerative Medicine, Strazhesko National Scientific Center of Cardiology, Clinical and Regenerative Medicine, 01024, Kyiv, Ukraine; Lund Stem Cell Center, Lund University, SE-22184, Lund, Sweden; Division of Molecular Hematology. Wallenberg Center for Molecular Medicine. Lund Stem Cell Center, Lund University, SE-22184, Lund, Sweden; Division of Neurosurgery, Department of Clinical Sciences Lund, University Hospital, SE-22184, Lund, Sweden

## Abstract

Neuronal loss and axonal demyelination underlie long-term functional impairments in patients affected by brain disorders such as ischemic stroke. Stem cell-based approaches reconstructing and remyelinating brain neural circuitry, leading to recovery, are highly warranted. Here we demonstrate the *in vitro* and *in vivo* production of myelinating oligodendrocytes from a human induced pluripotent stem (iPS) cell-derived long-term neuroepithelial stem (lt-NES) cell line, which also gives rise to neurons with the capacity to integrate into stroke-injured, adult rat cortical networks. Most importantly, the generated oligodendrocytes survive and form myelin ensheathing human axons in the host tissue after grafting onto adult human cortical organotypic cultures. This lt-NES cell line is the first human stem cell source that after intracerebral delivery can repair both injured neural circuitries and demyelinated axons. Our findings provide supportive evidence for the potential future use of human iPS cell-derived cell lines to promote effective clinical recovery following brain injuries.

## INTRODUCTION

Several disorders affecting the human brain, such as ischemic stroke, head trauma, and multiple sclerosis, lead to both neuronal loss and axonal demyelination, which underlie the functional deficits (Benedict et al., 2020; Kuhn et al., 2019; Nasrabady et al., 2018). A major cause of demyelination is the death of oligodendrocytes, i.e., the cell population producing myelin. The brain reacts to demyelination by two compensatory mechanisms: first, through the production of new myelin from oligodendrocytes in the area surrounding the lesion (Duncan et al., 2018), resulting in fewer and, in some cases, mistargeted myelin sheets (Neely et al., 2022); Second, by increased endogenous oligodendrogenesis (Franklin et al., 2021). However, this response is limited and most newly-generated oligodendrocytes fail to differentiate and remyelinate (Hughes et al., 2018; Marin and Carmichael, 2019).

Transplantation of remyelinating cells derived from stem cells has the potential to become a novel approach for treating myelin loss in the human brain. Oligodendrocytes can be generated from human induced pluripotent stem (iPS) cells or embryonic stem (ES) cells using different reprogramming and differentiation protocols (Ehrlich et al., 2017; Garcia-Leon et al., 2018; Shaker et al., 2021; Wang et al., 2013). Moreover, grafted oligodendrocytes produced from human pluripotent stem cells are capable of remyelinating host axons in the rodent’s central nervous system. Human ES cell-derived oligodendrocyte progenitor cells (OPCs), transplanted into the demyelinated mouse spinal cord and irradiated rat brain, remyelinated host-derived demyelinated axons (Nistor et al., 2005; Piao et al., 2015). Also, human iPS cell-derived OPCs and isolated O4^+^ population (containing immature and mature oligodendrocytes), grafted into mouse injured spinal cord and demyelinated corpus callosum, remyelinated axons (Ehrlich et al., 2017; Kawabata et al., 2016). Besides replacing myelinating cells, human stem cell-derived transplants can promote remyelination in animal models by other mechanisms. For example, intracerebral transplantation of human astrocytic-fated iPS cell-derived progenitors increased endogenous oligodendrogenesis and remyelination via the release of growth factors in mice with white matter stroke (Llorente et al., 2021). Whether human oligodendroglial progenitors, derived from iPS or ES cells, can remyelinate demyelinated tissue after transplantation in the adult human brain is unknown.

Effective repair in brain disorders using cell transplantation will require the replacement of both neurons and oligodendrocytes, capable of remyelinating host and grafted neurons. Currently, little is known whether the same human stem cell source can give rise to both functional neurons and myelinating oligodendrocytes after transplantation into the adult brain (Baker et al., 2017; Nori et al., 2011). We recently showed that human iPS cell-derived longterm neuroepithelial stem (lt-NES) cells, fated towards a cortical neuronal phenotype and transplanted into the rat cortex adjacent to an ischemic lesion, gave rise to functional cortical neurons. These neurons sent projections to both hemispheres, became integrated into host neural circuitry, and reversed sensorimotor deficits (Palma-Tortosa et al., 2020; Tornero et al., 2017). We also found that 40% of cells in the transplant, a substantial number of graft-derived cells in the corpus callosum, and a few of them in the thalamus and striatum, expressed the oligodendrocyte marker Sox10. Moreover, human-derived myelin basic protein (MBP) was observed close to the transplant, providing suggestive evidence that the graft-derived Sox10^+^ cells could be oligodendrocytes contributing to remyelination.

Here we demonstrate that the human iPS cell-derived lt-NES cells, primed towards a cortical neuronal phenotype, produce *bona fide* oligodendrocytes both *in vitro* and *in vivo.* The generated cells display the structural, molecular, and functional characteristics of human mature oligodendrocytes and myelinate lt-NES cell-derived axons in culture as well as host-derived axons after xenotransplantation into rat stroke-injured cortex and allotransplantation into human adult cortical organotypic cultures.

## RESULTS

### Cortically fated human lt-NES cells form myelinating oligodendrocytes in cell culture

In our previous study (Palma-Tortosa et al., 2020), we obtained some evidence for the presence not only of neurons but also of myelin-forming oligodendrocytes in the grafts after intracerebral transplantation of cortically primed human lt-NES cells in a rat stroke model. Here we wanted to determine, first *in vitro*, whether the lt-NES cells can form genuine functional oligodendrocytes in addition to neurons. We, therefore, differentiated lt-NES cells according to our cortical priming protocol for up to 21 days in the cell culture (Tornero et al., 2013), and analyzed protein and gene expression of different oligodendrocytes and neuronal markers.

Using immunocytochemistry, we found that undifferentiated lt-NES cells (at day 0) were positive for Sox10, a marker known to be expressed in both neuroectodermal cells and oligodendrocytes, but did not express the neuroblast marker doublecortin (DCX) or the pan-oligodendrocyte marker Olig2 (**Figure S1A**). While Sox10 expression decreased with time, the expression of DCX and Olig2 started following 4 days of differentiation with a tendency to increase (**Figure S1A**). Expression of MBP started on day 8 in cells displaying the bipolar morphology typical of OPCs. At later time points, we observed an increase in morphological complexity (branching) of MBP-expressing cells, characteristic of pre-myelinating and myelinating oligodendrocytes (Figure S1A). Flow cytometry showed increased O4 (a marker for pre-myelinating and myelinating oligodendrocytes) expression over time (**Figure S1D**). Similarly, using RT-qPCR, we observed that our cortical priming protocol gave rise to increased DCX, Olig2, and MBP gene expression starting at day 8 (**Figure S1B**).

We then asked if the cortically primed human lt-NES cell-derived oligodendrocytes, differentiated for 21 days, can myelinate human lt-NES cell-derived axons. At this time point, immunocytochemistry demonstrated the presence of 5% Olig2- and 3% CC1 (a marker of mature oligodendrocytes)-expressing cells in the cultures (**Figure 1A-B**). To verify that the observed CC1 expression was not due to the presence of astrocytes in the culture, we performed double-staining of CC1 with the astrocytic markers GFAP and S100ß. No colocalization was found (data not shown). Moreover, flow cytometry showed that 1.5% of the cells expressed O4 (**Figure 1C**, 1.5 ± 0.28%, n=4). Importantly, we found MBP expression adjacent to the axonal marker Neurofilament, suggesting that the oligodendrocytes in the culture could be involved in axonal myelination. (**Figure 1D**).

**Figure 1.**
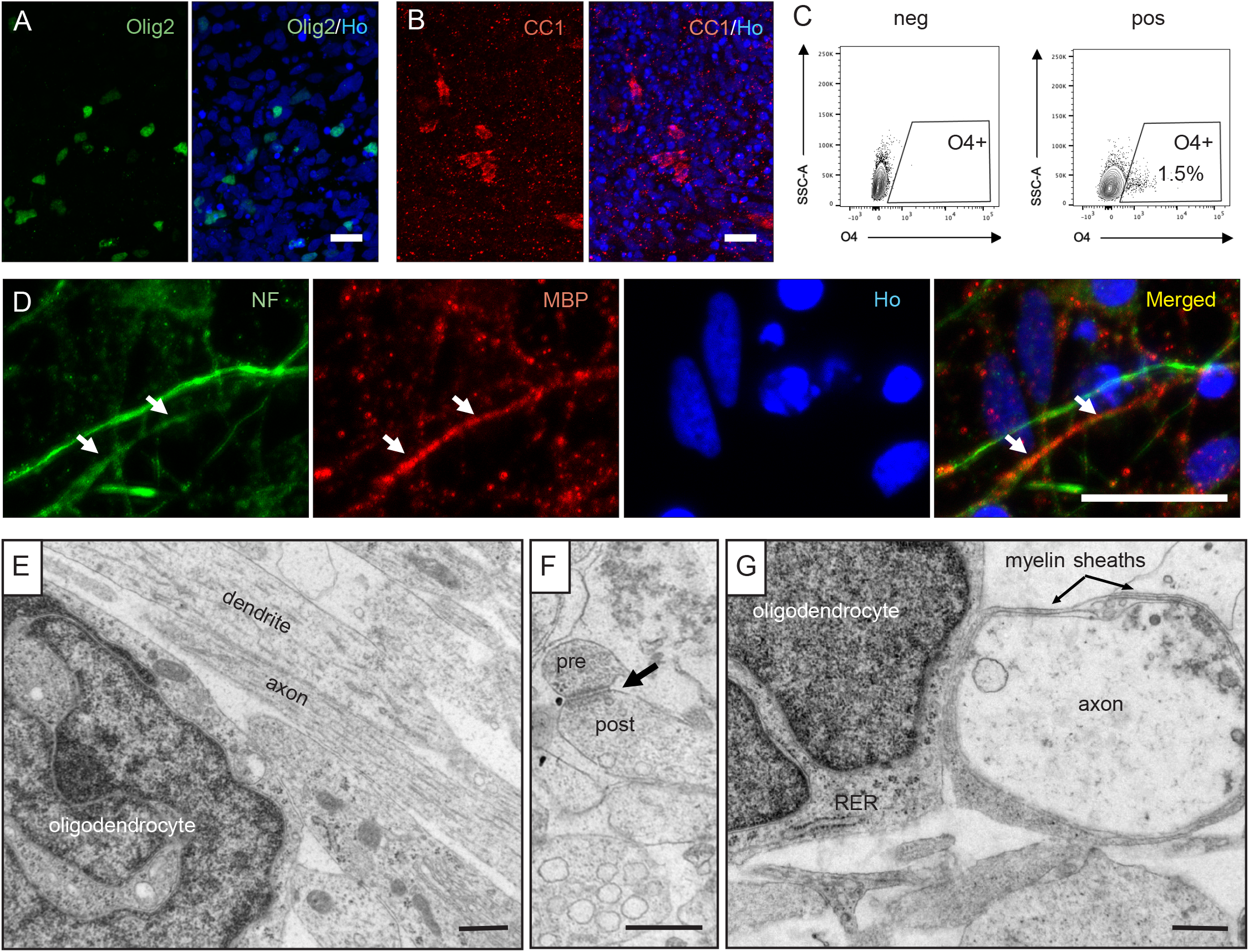
Cortically-primed human lt-NES cells generate mature neurons and myelinating oligodendrocytes after 21 days of *in vitro* differentiation. (**A**-**B**) Confocal images of lt-NES cells differentiated for 21 days showing the expression of (**A**) the pan-oligodendrocyte marker Olig2 and (**B**) the mature oligodendrocyte marker CC1. (**C**) Flow cytometry analysis of O4^+^ cells, including pre-myelinating and myelinating oligodendrocytes, after 21 days of differentiation (n=4 replicates). (**D**) Confocal immunohistochemical images showing axons immunoreactive for Neurofilament and myelin basic protein (MBP) expressed in closed proximity in culture (arrows depict proximity between both markers). Ho, Hoechst. Scale bar in A-D, 20 μm. (**E-G**) EM images showing: (**E**) a mature lt-NES cell-derived oligodendrocyte and neural processes, (**F**) an axodendritic contact with structural characteristics of the asymmetric synapse (arrow indicates a synapsis containing pre and post-synaptic components), and (**G**) lt-NES cell-derived myelin loosely wrapped around an lt-NES cell-derived axon. RER, rough endoplasmic reticulum. Scale bar in E-G, 500 nm.

We used electron microscopy (EM) to provide further evidence for the presence of lt-NES cell-derived oligodendrocytes and their ability to myelinate human axons in the cultures. In addition to neurons (Gronning Hansen et al., 2020), we detected a cell population exhibiting the ultrastructural features of mature oligodendrocytes, i.e., irregularly shaped dark nucleus with clumped chromatin near the inner nuclear membrane (**Figure 1E**). The cytoplasm was electron-dense and contained short cisternae of rough endoplasmic reticulum with short mitochondria. A large number of dendrites and axons was observed in the neuropil (**Figure 1E**). The neural processes formed axodendritic contacts with structural characteristics of asymmetric synapses (**Figure 1F**), suggesting that the lt-NES cell-derived neurons had established a network already at 21 days in culture. Importantly, the lt-NES cell-derived oligodendrocytes formed loose myelin sheaths around axons, which could be the initial stage of myelination (**Figure 1G**).

### Cortically fated human lt-NES cells form oligodendrocytes and myelinate host axons after transplantation in stroke-injured rat cortex

We then explored whether intracerebrally transplanted, cortically fated lt-NES cell-derived progenitors could remyelinate axons after stroke, i.e., an injury-causing axonal demyelination (Zuo et al., 2019). Rats were subjected to cortical stroke, implanted with human lt-NES cells close to the injury after 48 h, and sacrificed 6 months later. In all the animals, the stroke-induced neuronal loss, as determined by lack of NeuN (marker of mature neurons) immunoreactivity, was restricted to the cortex (mostly somatosensory cortex (S1FF and S1BL) and motor area (M1)), sparing subcortical structures (Palma-Tortosa et al., 2020).

To study the overall distribution of axonal demyelination following stroke, we quantified the expression of MBP as a measure of myelin density (**Figure 2**). The ischemic insult caused decreased MBP expression in the dorsal striatum, the middle part of the corpus callosum and its thickness, and peri-infarct areas (**Figure 2A-E, G**). Ectopic MBP expression was detected around Olig2^+^ cell bodies close to the injury in stroke animals (**Figure 2H-I**). Transplantation of lt-NES cells resulted in increased MBP expression in the corpus callosum and dorsal striatum and a similar tendency in the peri-infarct area compared to non-grafted rats, whereas the thickness of corpus callosum did not differ between the groups of stroke-affected animals (**Figure 2A-G**). The presence of lt-NES cells in areas of demyelination was mainly restricted to the peri-infarct zone and corpus callosum, illustrating that the contribution of grafted cells to remyelination was limited and indicating that other myelination mechanisms were most likely triggered by the transplantation (**Figure 2A-D**).

**Figure 2.**
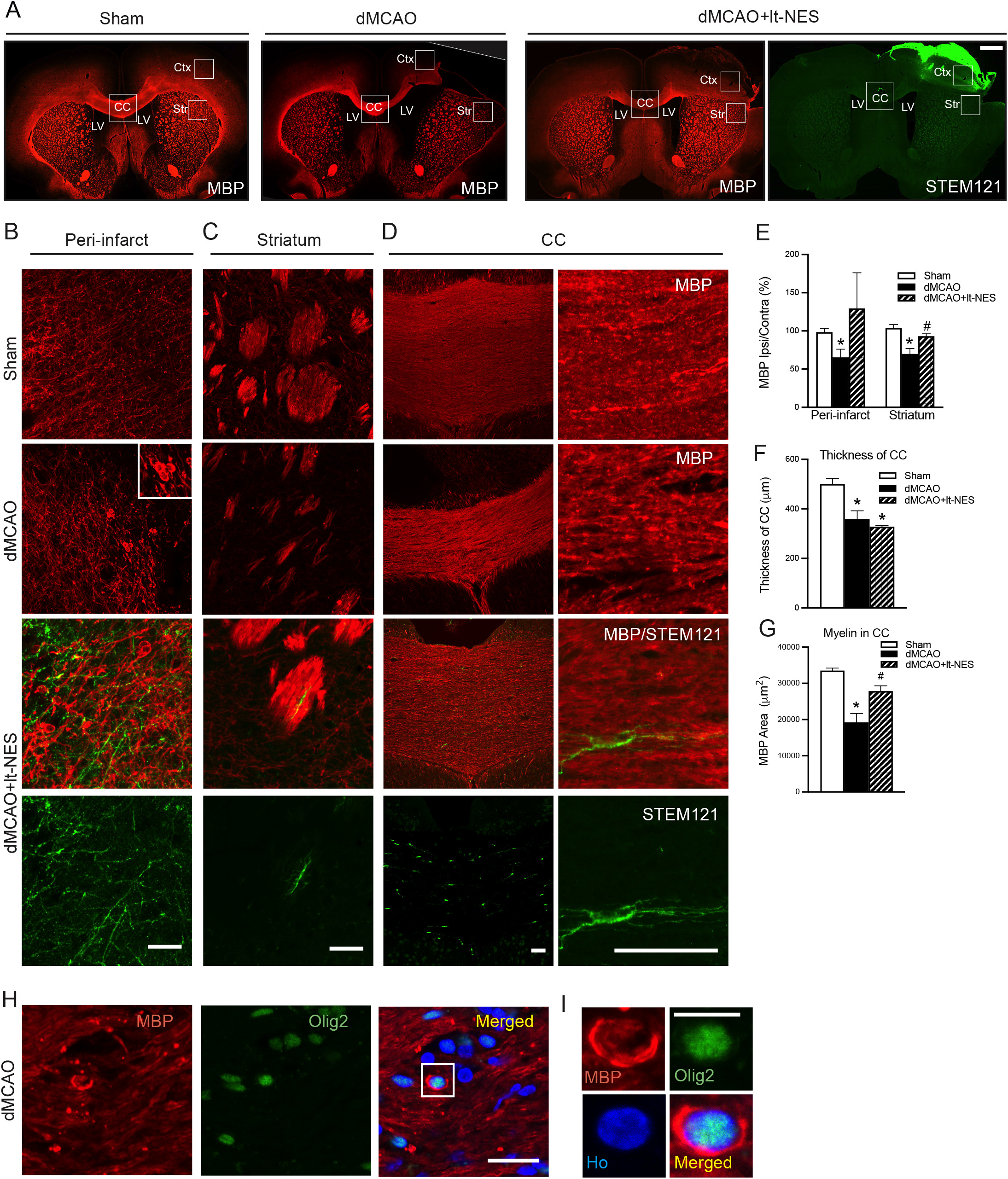
Intracortical transplantation of human lt-NES cell-derived progenitors improves myelination in stroke-injured rat brains. (A) Coronal overview of MBP^+^ area in the rat brain of sham-treated animals (Sham), and in animals subjected only to stroke (dMCAO) or stroke followed by transplantation (dMCAO+lt-NES). An overview of the area positive for the human cell cytoplasmic marker STEM121^+^ is also included in the transplanted group. White squares show the areas where images were taken and quantification has been performed. Scale bar in A, 1 mm. CC: Corpus callosum; Ctx: Cortex; LV: Lateral ventricle; Str: Striatum. (B-G) Confocal immunohistochemical images and quantification of MBP^+^ area in (B, E) periinfarct area, (C, E) dorsal striatum, and (D, F, G) middle part of the corpus callosum of Sham, dMCAO or dMCAO+lt-NES. STEM121 staining shows the contribution of graft-derived cells in different areas. Scale bar in B-D, 100 μm. n=4-5 animals per group. (H-I) Image showing myelin surrounding Olig2^+^ cell (higher magnification in H). Nuclear staining (Hoechst, blue) is included in the merged panel. Scale bar in H-I, 20 μm. *p<0.05 vs. Sham. #p<0.05 vs. dMCAO. Data are shown as mean ± SEM.

Our previous study indicated that the grafted lt-NES cell-derived neurons regulate motor behaviour in stroke-injured rats through transcallosal connections to the contralateral hemisphere (Palma-Tortosa et al., 2020). We therefore analyzed in more detail the effect of stroke and lt-NES cell transplantation on the number of oligodendrocytes in the corpus callosum. Immunohistochemical analysis showed more Olig2^+^ cells in the middle part of the corpus callosum in animals subjected to stroke compared to sham-treated animals. In transplanted rats, the increase of Olig2^+^ cells was even more pronounced. The majority of Olig2^+^ cells did not co-express the human nuclear marker STEM101 and were most likely rat-derived cells. Some human cells were also found, even if they contributed only to a fraction of the total number of Olig2^+^ cells (**Figure S2A-C**).

Since we detected Olig2^+^ cells of human origin in the corpus callosum, we hypothesized that part of the transplanted lt-NES cell-derived progenitors had become oligodendrocytes and contributed to the remyelination. Arguing for the formation of mature oligodendrocytes *in vivo,* we found that at 6 months after transplantation, 15-20% of the STEM101^+^ cells in the core of the graft expressed Olig2 (**Figure 3A**) and about 20% expressed CNPase (marker for pre-myelinating and myelinating oligodendrocytes) (**Figure 3C**). We also observed human-derived cells expressing the mature oligodendrocyte marker, CC1, with varying density through the graft (**Figure 3D**). Hardly any grafted cells expressed neuron-glial antigen 2 (NG2) (**Figure 3B**), a marker for OPCs.

**Figure 3.**
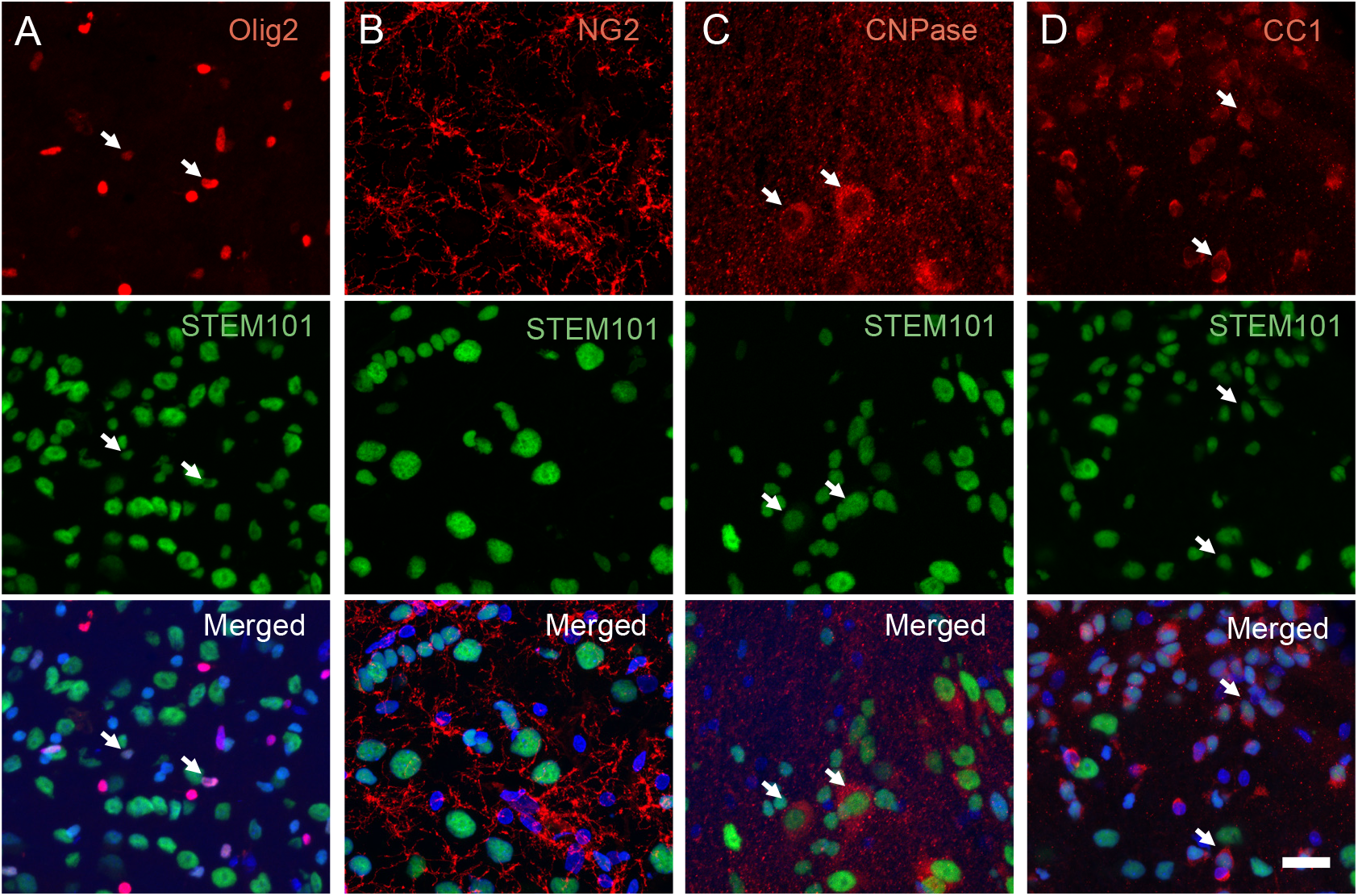
Cortically-primed human lt-NES cell-derived progenitors give rise to mature oligodendrocytes 6 months after intracortical transplantation into stroke-injured rat somatosensory cortex. Representative confocal images of the transplantation area showing the expression of (**A**) the pan-oligodendrocyte marker Olig2, (**B**) OPC marker NG2, (**C**) marker for immature and mature oligodendrocytes CNPase, and (**D**) mature oligodendrocyte marker CC1. Arrows indicate the colocalization of respective markers with STEM101, a human nuclear marker. Nuclear staining (Hoechst, blue) is included in the merged panel. Scale bar, 20 μm.

To provide further evidence for the formation of mature myelinating oligodendrocytes, we performed the ultrastructural analysis 6 months after transplantation in stroke-affected animals with grafts of GFP-labeled, cortically fated human lt-NES cells. Following the validation of GFP expression in myelin sheets in our cell line (data not shown), immunoperoxidase reaction or immunogold staining using anti-GFP antibodies was carried out to localize the grafted GFP^+^ lt-NES cells and their processes, which were easily identified due to the brown DAB reaction product in cytoplasm and processes (**Figure 4A-B**). Most GFP^+^ cells were located in the peri-infarct area while some of them were found in the corpus callosum and contralateral cortex. Some GFP^+^ cells exhibited the morphology of mature myelinating oligodendrocytes (**Figure 4C**): dark electron-dense rectangular-shape cytoplasm, heterogeneous nuclear chromatin pattern, and short and wide endoplasmic reticulum cisternae organized in the vicinity of the nucleus.

**Figure 4.**
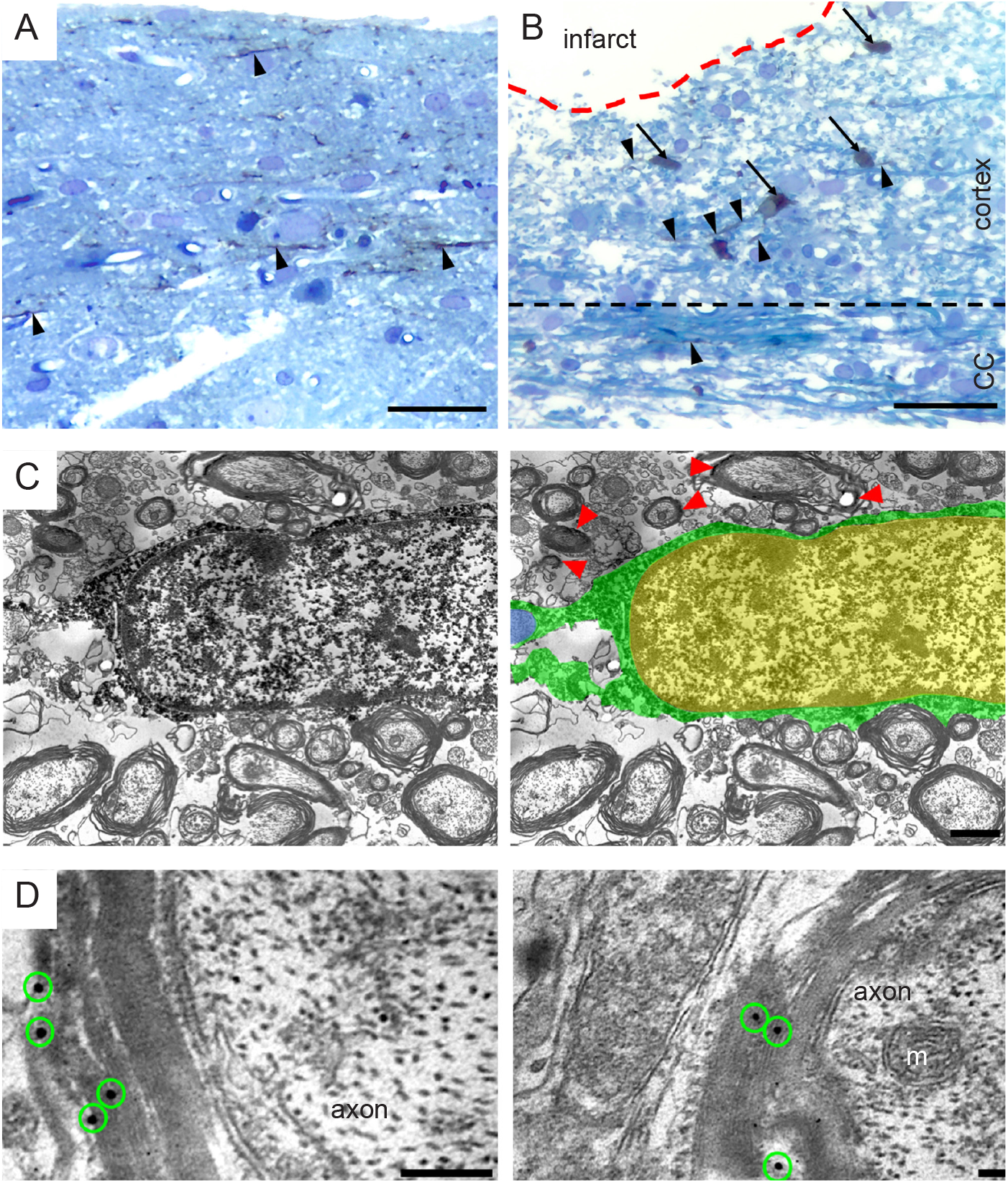
Grafted human lt-NES cell-derived oligodendrocytes myelinate axons in the rat injured cortex. (**A**-**B**) Light micrographs of a toluidine blue-stained plastic-embedded section from the cortex of stroke subjected rats contralateral (**A**) and ipsilateral (**B**) to the lesion. GFP/DAB-positive lt-NES cell-derived cells were visualized in the peri-infarct area (ipsilateral cortex), corpus callosum, and contralateral cortex. Arrows highlight GFP/DAB-positive cells (brown color), and arrowheads indicate GFP/DAB-positive processes (brown color). (**C**) Representative immunoelectron microscopy (iEM) image showing that at 6 months after transplantation, GFP/DAB-positive cells in the corpus callosum exhibit the morphology of mature myelinating oligodendrocytes (for better visualization in the electron microscopical images, GFP/DAB-positive oligodendrocyte cytoplasm and processes are colored green and the nucleus is yellow). The host axon myelinated by the GFP/DAB-positive oligodendrocyte is colored blue. Red arrowheads indicate areas of GFP/DAB-positive myelin. (**D**) Electron micrographs of GFP^+^ immunogold particles (green circles) associated with compact myelin sheaths. CC: corpus callosum. m, mitochondrion. Scale bar in A-B, 50 μm. Scale bar in C, 2 μm. Scale bar in D, 0.1 μm.

To determine if the human lt-NES cell-derived oligodendrocytes could form myelin sheaths around host axons, we performed post-embedding immunogold labeling of GFP. In support of host axonal myelination by the graft-derived human oligodendrocytes, iEM demonstrated individual or clusters of gold particles within the membranous sheets of myelin (i.e., graft-derived myelin) surrounding host unlabeled axons (**Figure 4D**).

### Grafted human lt-NES cell-derived oligodendrocytes myelinate host axons in adult human cortical tissue

We have previously shown that cortically fated lt-NES cell-derived progenitors differentiate to cortical neurons and establish functional connections with host neurons after transplantation onto organotypic cultures of adult human cortex obtained from epileptic patients undergoing surgery (Gronning Hansen et al., 2020). Here we used this model to determine whether grafted lt-NES cells also give rise to mature oligodendrocytes which can myelinate axons in the adult human cortical environment.

We first assessed the preservation of the cortical tissue during the organotypic culture. In accordance with our previous findings (Gronning Hansen et al., 2020; Palma-Tortosa et al., 2020), we found that the expression of the neuronal marker NeuN was visually unaffected after 4 weeks in culture compared to acute tissue, whereas Map2 expression had decreased by more than 90% (**Figure 5A-B**). Olig2, CNPase, and CC1 were expressed in acute tissue (**Figure S3A-B**), and MBP expression resembled the typical aligned axonal distribution in the adult human cortex (**Figure S3A**). After 4 weeks in culture, the expression of Olig2, CNPase, and CC1 was unaffected (**Figure S4A-C**). Overall MBP expression was diminished compared to acute tissue **(Figure S3A**).

**Figure 5.**
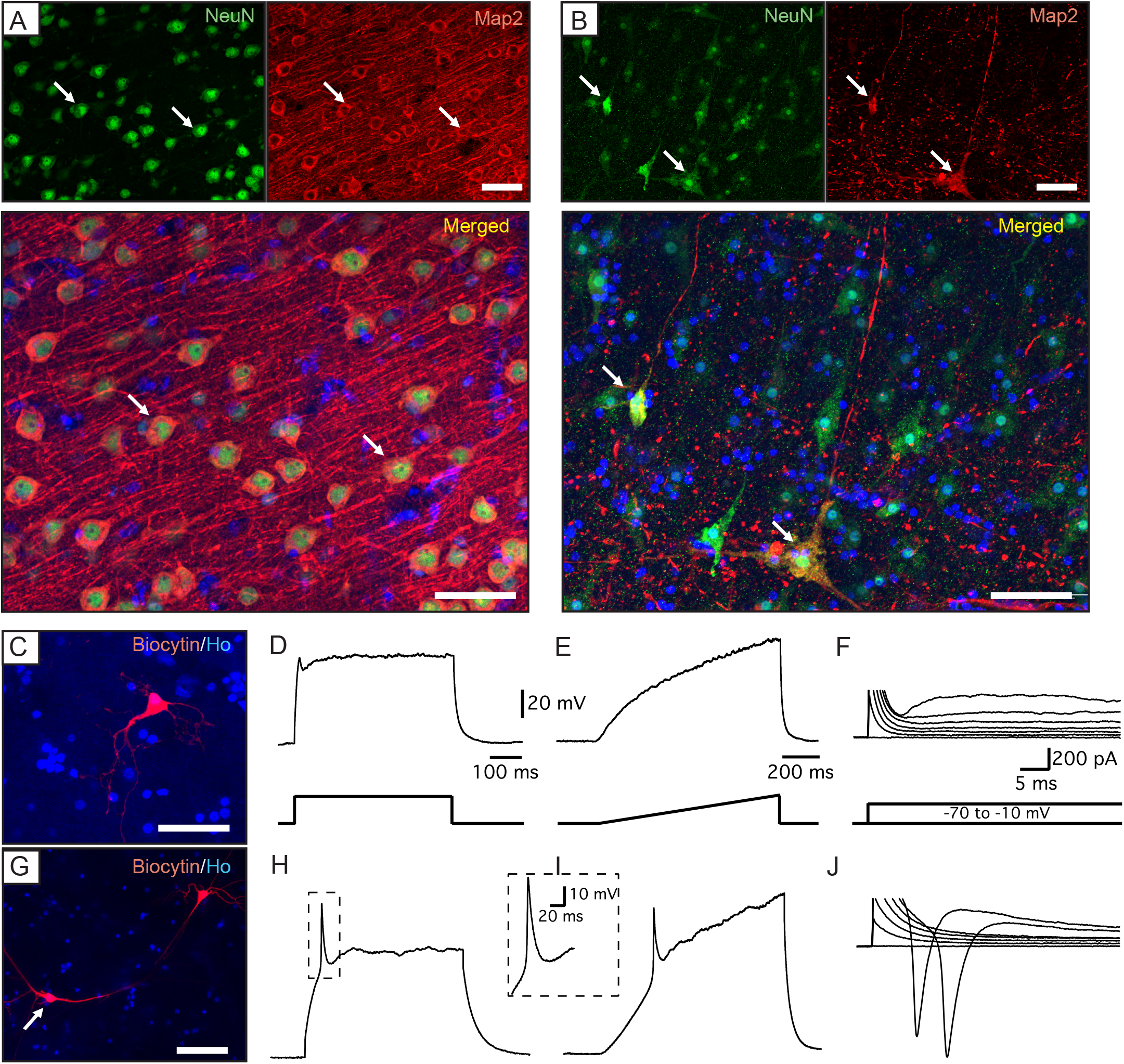
The functionality of neuronal circuitry is compromised in human adult cortical organotypic tissue after 4 weeks in culture. (**A-B**) Representative confocal images of (**A**) acute and (**B**) 4 weeks-cultured human cortical tissue showing the expression of the neuronal markers NeuN and Map2. Arrows indicate colocalization. Nuclear staining (Hoechst, blue) is included in the merged panel. (**C-J**). Examples of cortical neurons recorded in 4 weeks-cultured human cortical tissue. (**C**, **G**) Biocytin labelling of the recorded neurons. (**D**-**F**) Whole-cell patch-clamp recording traces from the cell in (**C**). The cell is not able to fire AP either at step (250 pA) (**D**) or at the ramp (0 to 300 pA) (**E**) current injection in current clamp mode; (**F**) the same cell has a very small inward sodium current upon 10 mV depolarization step in voltage-clamp mode at −70 mV holding potential. (**H-J**) Patch-clamp recording of another cell, shown in (**G**), that was able to generate a single AP in the similar protocol as above (**H**) and (**I**), with a bigger sodium current in voltage-clamp mode (**J**). Scale bars are the same for the upper and lower panels of the traces unless otherwise indicated. Ho, Hoechst. Scale bar, 50 μm

Electrophysiological analysis showed that 2 out of 7 recorded neurons were able to fire a single action potential (AP) (**Figure 5G-I**), while the majority were unable to generate any (5 out of 7) (**Figure 5C-E**). Inability to fire AP corresponded with small inward sodium currents, indicating that sodium conductance, which is essential for normal neuronal function and an initial phase of AP, was compromised (**Figure 5F, J**). However, passive membrane properties, such as membrane resistance (301 ± 67 MΩ) and resting membrane potential (−74,7 ± 2,5 mV), were still comparable to the values reported in fresh adult human cortical tissue (Avaliani et al., 2014), indicating that the neurons are still alive and relatively viable, although not very active.

Taken together, our data indicate that the architecture of the adult human cortical tissue is largely preserved but the functionality of the neuronal network is decreased after 4 weeks in culture.

To assess their capacity to differentiate into myelinating oligodendrocytes in the adult human cortical environment, cortically fated GFP^+^ lt-NES cells were then transplanted onto the organotypic slice cultures (**Figure S5**). We found graft-derived (GFP^+^ cells) with extensive arborizations throughout the organotypic culture after 4 weeks (**Figure 6A**). About 40-50% of the GFP^+^ cells colocalized with the pan-oligodendrocyte marker Olig2 (**Figure 6B-C**). Importantly, colocalization of GFP^+^ cells with either CNPase or CC1 were also found, arguing for the presence of both immature and mature graft-derived oligodendrocytes (**Figure 6D-G**). Due to the cytoplasmic nature of CNPase and CC1 staining and since a human cell marker cannot be used to identify human cells transplanted into human tissue, the percentage of grafted cells positive for these markers could not be evaluated.

**Figure 6.**
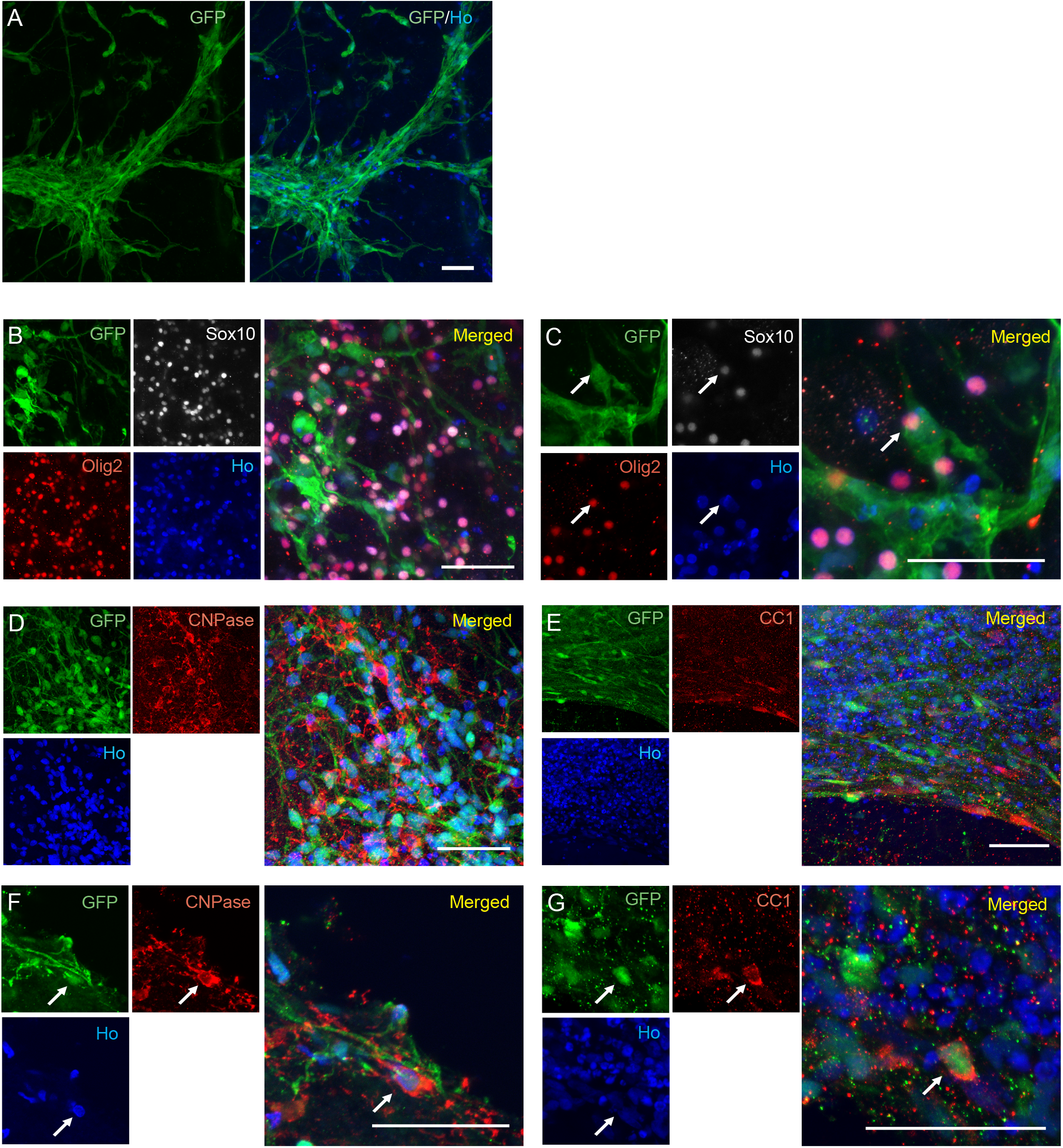
Human lt-NES cell-derived progenitors survive long-term and become mature oligodendrocytes 4 weeks after *ex vivo* transplantation into organotypic cultures of the human adult cortex. (**A**) Overview of grafted GFP^+^ lt-NES cells 4 weeks after transplantation into the human cortical culture. (**B**-**G**) Confocal images showing GFP^+^ lt-NES cells after *ex vivo* transplantation expressing (**B**-**C**) Sox10 and Olig2, (**D, F**) CNPase, and (**E, G**) CC1. (**B, D, and E**) Low magnification images. (**C, F, and G**) High magnification images. Ho, Hoechst. Arrows indicate colocalization. Scale bar, 50 μm.

We finally determined if the grafted human GFP^+^ lt-NES cell-derived oligodendrocytes had formed myelin sheaths. Anti-GFP antibodies and immunogold labeling combined with iEM demonstrated the presence of individual or clusters of gold particles within the membranous sheets of myelin (signifying graft-derived myelin) surrounding non-GFP-labeled host axons (**Figure 7A-B**). These results provide strong evidence that lt-NES cell-derived oligodendrocytes can myelinate human-derived axons after transplantation into adult human cortical tissue.

**Figure 7.**
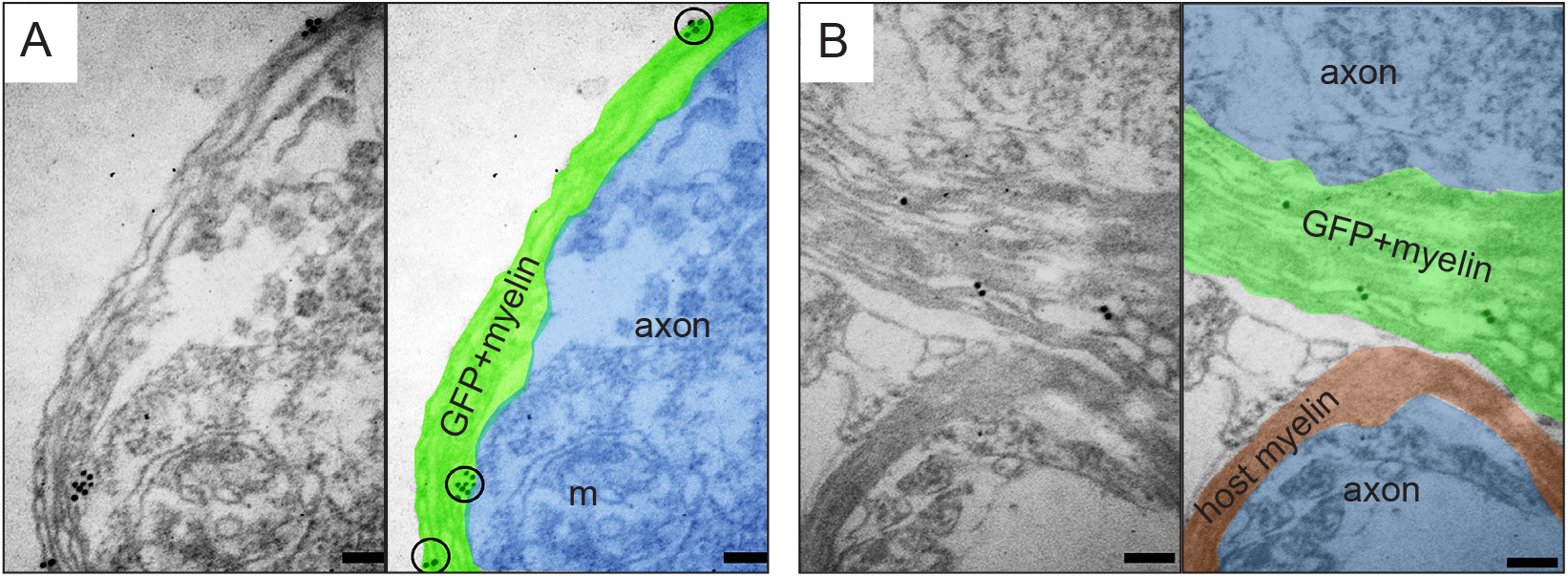
Human lt-NES cell-derived oligodendrocytes myelinate human-derived axons 4 weeks after *ex vivo* transplantation into organotypic cultures of the human adult cortex. (**A**-**B**) iEM images showing human grafted GFP^+^ lt-NES cell-derived myelin wrapping human axons in organotypic cultures. For better visualization in the electron microscopical images, GFP/gold-positive myelin is colored in green, GFP-negative myelin is colored in red, and axons in blue. Black circles depict immunogold particles associated with compact myelin sheaths. m, mitochondrion. Scale bar, 0.1 μm.

## DISCUSSION

Here we describe the *in vitro* and *in vivo* production of myelinating oligodendrocytes from a human iPS cell-derived cell line which also gives rise to neurons with the capacity to integrate into the adult human neural circuitry (Palma-Tortosa et al., 2020). Most importantly, the generated oligodendrocytes survive and form myelin ensheathing human axons in the host tissue after grafting onto adult human cortical organotypic cultures. We have previously shown that the human iPS cell-derived cortical neurons establish connections with the human host neurons in these slice cultures (Gronning Hansen et al., 2020). Taken together, we report for the first time a human iPS cell-derived cell line, primed towards a cortical neuronal phenotype, with the capacity to generate both neurons and oligodendrocytes reconstructing neural circuitry and myelinating axons in the adult human cerebral cortex.

Our conclusion that the generated cells are *bona fide* functional oligodendrocytes is inferred from the following findings: First, the lt-NES cell-derived progenitors expressed oligodendrocyte lineage markers, such as Olig2, O4, CNPase, and CC1, as evidenced by immunocytochemistry and flow cytometry; Second, EM showed the presence of mature oligodendrocytes and myelin around lt-NES cell-derived axons after 21 days of *in vitro* differentiation; Third, 6 months after intracortical transplantation into stroke-injured rat cortex, 20% of grafted cells had become oligodendrocytes, expressing Olig2, CNPase, and CC1, and graft-derived myelin was present in areas of demyelination, as observed with MBP immunohistochemistry and EM; Fourth, 1 month after transplantation onto adult human cortical cultures, grafted cells expressed markers for mature oligodendrocytes (CNPase and CC1), and graft-derived myelin was found around host human axons using immunohistochemistry and EM.

Generally, human neural progenitor cell- or stem cell-derived OPCs are used to generate myelinating oligodendrocytes (Chanoumidou et al., 2020) and neural differentiation protocols to produce functional neurons (Tornero et al., 2013; Zhao et al., 2020). Rapid concomitant generation of both mature neurons and myelinating oligodendrocytes from human stem cells in culture is a challenge. For example, differentiation of human iPS cell-derived lt-NES cells into both neuronal and glial lineages gave rise to mature neurons but, even if oligodendrocytic markers were expressed, no oligodendrocyte-like morphology or axonal myelination was identified after 4 weeks (Isoda et al., 2016). Oligodendrocyte generation in a neuronally committed culture can be improved by the addition of factors that promote glial proliferation and differentiation. Thus, Ehrlich and co-workers (Ehrlich et al., 2017) reported that overexpression of Sox10, Olig2, and NKX6.2 in human iPS cell-derived neural progenitors, combined with platelet-derived growth factor (PDGF-AA), smoothened agonist (SAG) and thyroid hormone (T3), accelerates oligodendroglial differentiation, reaching 70% oligodendrocytes and 20% neurons after 28 days *in vitro*. Importantly, MBP expression was observed at day 35 of the differentiation (Ehrlich et al., 2017). Shaker and collaborators have shown that the generation of organoids from human neuroectodermal cells in the presence of T3, neurotrophin 3, hepatocyte growth factor (HGF), insulin growth factor (IGF), PDGF-AA, and cAMP gives rise to 29% neurons, 20% immature oligodendrocytes and 40% myelinating oligodendrocytes at day 42 of differentiation (Shaker et al., 2021). We found here a relatively low percentage of oligodendrocytes (5%) using our cortical differentiation protocol *in vitro.* However, functional myelination was observed already after 21 days of differentiation, i.e., earlier as compared to previous protocols (Ehrlich et al., 2017; Shaker et al., 2021). It seems possible that also in our cultures, increased numbers of oligodendrocytes could be obtained using some of the factors mentioned above.

Previous studies with human pluripotent stem cells have reported the production of either neurons or oligodendrocytes after transplantation. In grafts of human iPS cell-derived O4^+^ cells in the spinal cord of demyelinating mice, around 80% of the cells were oligodendrocytes and only 2% expressed neuronal markers (Ehrlich et al., 2017). Similarly, human ES cell-derived OPCs, transplanted in spinal cord-injured rats gave rise to 94% oligodendrocytes (Sharp et al., 2010). Conversely, other intracerebral ES or iPS cell-derived grafts have mainly contained neurons. At 60 weeks after transplantation of human ES cell-derived lt-NES cells in mouse dentate gyrus and motor cortex, about 80% of cells in the grafts were neurons and no oligodendrocytes were found (Steinbeck et al., 2012). Likewise, transplantation of iPS cell-derived lt-NES cells in spinal cord injured mice gave rise to 75% mature neurons and less than 1% oligodendrocytes after 7 weeks (Fujimoto et al., 2012).

Whether substantial numbers of both functional neurons and myelinating oligodendrocytes can be produced from the same intracerebral human stem cell grafts is not well studied. Nori and collaborators found a majority of neurons and 9% oligodendrocytes in grafts of human iPS cell-derived neurospheres at 112 days after transplantation in spinal cord-injured mice (Nori et al., 2011). Intracortical transplantation of human iPS cell-derived neural stem cells in a pig model of cortical stroke gave rise to 75% neurons and 25% oligodendrocytes (Baker et al., 2017). These percentages are consistent with our observations of 40% mature neurons and 20% pre-myelinating and myelinating oligodendrocytes after intracortical transplantation of cortically primed lt-NES cells into stroke-injured rat cortex. Compared to the studies of Nori and co-workers (Nori et al., 2011) and Baker and co-workers (Baker et al., 2017), we show here for the first time that grafted human pluripotent stem cell-derived oligodendrocytes can be functional and myelinate axons. Most importantly, we demonstrate the survival of a large number of these oligodendrocytes and their capacity for myelination also after transplantation in an *ex vivo* model of adult human cerebral cortex. It should be pointed out, though, that our study was only designed to show proof-of-principle that grafted human iPS cell-derived oligodendrocytes can myelinate human axons. Further analysis needs to be done to determine the extent of this myelination in the human system.

We found increased endogenous oligodendrogenesis in response to stroke as reported previously (Hernandez et al., 2021). Transplantation of human iPS cell-derived progenitors further enhanced endogenous oligodendrogenesis and increased axonal remyelination in several demyelinated brain areas. Such potentiation of oligodendrogenesis, attributed to a so-called bystander effect, has been observed also following administration of other human stem cell types, e. g., after intracerebral transplantation of human iPS cell-derived immature astroglia in a mouse model of neonatal white matter injury (Jiang et al., 2016) or intravenous delivery of human adipose-derived stem cells in a rat demyelinated model (Bakhtiari et al., 2021). Only a minor portion of the new oligodendrocytes originated from the grafted, cortically primed human lt-NES cells. This finding indicates that the migration of graft-derived progenitors from the transplant core to demyelinated areas and the capacity to become myelinating oligodendrocytes need to be increased to optimize functional recovery. It is important to point out, though, that the low contribution to remyelination by the grafted human-derived oligodendrocytes and the dominant role of endogenous oligodendrogenesis observed here in the rat stroke model may not reflect the outcome in a hypothetical clinical setting. Since endogenous oligodendrogenesis and remyelination efficiency is decreased by aging (Segel et al., 2019; Shen et al., 2008), replacement by grafted oligodendrocytes would probably be more critical for remyelination in older patients, who are most often affected by stroke.

Current stem cell-based approaches which have reached patient application in stroke aim at stimulating plasticity and dampening inflammation (de Celis-Ruiz et al., 2021; Gong et al., 2021; Hess et al., 2017) but only minor or no improvements have been so far been observed. From a clinical perspective, a human cell source that after intracerebral delivery has the capacity both for the functional repair of injured neural circuitries and for remyelination could be a step forward toward effective stem cell therapy for patients not only with stroke but also other brain injuries. The present findings provide supportive evidence that the further development of such a cell source is a realistic possibility.

## EXPERIMENTAL PROCEDURES

### Resource availability

#### Corresponding author

Further information and requests for resources and reagents should be directed to and will be fulfilled by the corresponding author, Zaal Kokaia.

#### Materials availability

This study did not generate new unique reagents.

#### Data and code availability

The data that support the findings of this study are available on request from the corresponding author.

### Generation of iPS cell-derived oligodendrocytes and cortical neurons

Human iPS cell-derived lt-NES cells were produced as previously described (Falk et al., 2012; Gronning Hansen et al., 2020) (see **supplemental experimental procedures**). lt-NES cells used for transplantation onto *ex vivo* human tissue were transduced with a lentiviral vector carrying GFP under constitutive promoter (GFP^+^ lt-NES cells). Differentiation of lt-NES cells to oligodendrocytes and neurons with a cortical phenotype was performed as previously described (Gronning Hansen et al., 2020; Tornero et al., 2013). Briefly, growth factors (EGF, bFGF) and B27 were omitted and lt-NES cells were cultured at low density in a differentiation-defined medium (DDM) in the presence of bone morphogenetic protein 4 (BMP4) (10 ng/mL, R&D Systems), wingless-type MMTV integration site family, member 3A (Wnt3A) (10 ng/mL, R&D Systems) and cyclopamine (1 μM, Calbiochem) for 7 days. On day 7, media was changed to Brain Phys (StemCell Technologies) supplemented with B27 without retinoid acid (1:50, Invitrogen) until day 21 of differentiation.

### Animals and surgical procedures

All procedures were conducted following the European Union Directive (2010/63/EU) and were approved by the ethical committee for the use of laboratory animals at Lund University and the Swedish Board of Agriculture (Dnr. M68-16).

Adult athymic, nude male rats (220 g, n= 18; Charles River) were used (5 sham-treated animals, 5 stroke-subjected animals, and 8 for stroke and cell transplantation). Among transplanted animals, five were destinated for immunological characterization of the grafts, and 3 for immuno-electron microscopy (iEM). Details for cortical ischemic injury and cell transplantation in the somatosensory cortex are described in **supplemental experimental procedures.**

### Organotypic cultures of the human adult cortex

Healthy human neocortical tissue was obtained with informed consent by resection of a small piece of the middle temporal gyrus from patients undergoing elective surgery for temporal lobe epilepsy (n=3) according to guidelines approved by the Regional Ethical Committee, Lund (Dnr. 2021-07006-01). The tissue slices were delivered and handled as previously described (Gronning Hansen et al., 2020; Miskinyte et al., 2017). Details in the processing of the human slices are described in **supplemental experimental procedures.**

### Transplantation of lt-NES cells onto human organotypic cortical slices

The lt-NES cell transplantation was performed as previously described (Gronning Hansen et al., 2020). Briefly, GFP^+^ lt-NES cells were detached at day 7 of differentiation and resuspended at a concentration of 100000 cells/μL in pure cold Matrigel Matrix (Corning). After part of the media was removed from the top of the insert, the suspension mix was collected into a cold glass capillary and injected as small drops stabbing the semi-dry slice at various sites. After the matrigel was solidified, additional media were carefully added to completely immerse the tissue. The media were changed once a week and co-culture was maintained for 4-6 weeks before fixation.

### Immunostainings and quantifications

Details of the immunocytochemistry in cultured cells and immunohistochemistry in rat slices and human organotypic slices are described in **supplemental experimental procedures**.

Overview images of rat brain slices stained with Olig2 and MBP were taken using a Virtual Slide Scanning System (VS-120-S6-W, Olympus, Germany).

CNPase and Olig2 quantification in the core of the transplantation in the rat slices was performed using 40x confocal images (LSM 780, Zeiss, Germany) and numbers of positive cells were counted by sampling different areas of the core of the transplant ((CNPase^+^-STEM101^+^)/total STEM101^+^). Co-expression was assessed in the confocal microscope as overlapping of the two selected markers in the same plane and area.

Quantification of Olig2^+^ cells in the corpus callosum in the rat slices was performed using 20x confocal images (10 μm thick for Olig2 staining). For analysis of myelination, confocal images (5μm thick for MBP staining) were taken in the corpus callosum (40x), peri-infarct area, and striatum (20x). For analysis of the dorsal lateral striatum, starting 700 μm lateral to the dorsal part of the lateral ventricle, two 20x images (x= 700 μm and 1400 μm) were taken. To measure the thickness of the corpus callosum confocal images were taken with 20x Zoom 0.6.

Quantification of GFP^+^ Olig2^+^ cells in the human organotypic slices was carried out using 20x confocal images in areas where transplantation was observed, and number of double positive cells were counted through a Z-stack by sampling different single planes.

All quantifications were performed on maximum-intensity projection images using ImageJ software by blinded researchers.

### Flow cytometry

The lt-NES cell cultures were harvested and washed before antibody incubation. Anti-O4 APC-conjugated antibody (Miltenyi) was diluted 1:200 in FACS buffer (PBS + 2% fetal bovine serum [FBS] + 2 nM ethylenediaminetetraacetic acid [EDTA]). Cells were incubated for 30 min at +4°C in darkness, followed by wash and incubation with propidium iodide (Life Technologies), a viability marker, diluted 1:1000 in FACS buffer at least 5 min before acquisition. Cells were analyzed in an LSR II flow cytometer (Becton Dickinson) and results were analyzed with BD FACSDiva 9.0 software (BD Biosciences) and FlowJo™ v10.8 Software (BD Life Sciences). The gating strategy is shown in **Figure S1C**.

### Quantitative reverse transcription polymerase chain reaction (RT-qPCR)

RT-qPCR was performed on RNA extracted from cells at different time points of differentiation (D0, D4, D8, D12, and D15). RNA extraction was performed with the RNeasy Mini kit (Qiagen) following the protocol described by the manufacturer. RNA purity and concentration of samples were determined using a Nanodrop spectrophotometer (ND-1000). 1μg of RNA was used for cDNA synthesis with Qscript™ cDNA SuperMix (QuantaBio). TaqMan probes (ThermoFisher Scientific; Olig2, HS00300164_s1; MBP, HS00921945_m1; DCX, HS00167057_m1; GAPDH, HS02786624_g1) were used and RT-qPCR was run in triplicate samples on an iQ5 real-time cycler (Bio-Rad) with GAPDH as the housekeeping gene.

### Electron microscopy

For *in vitro* EM, lt-NES cell cultures (n=8 wells) were fixed with 2% formaldehyde and 0.2% glutaraldehyde in 0.1 M phosphate buffer, pH 7.4. Samples were frozen and cut into ultrathin sections with a diamond knife. Ultrathin sections were examined and photographed using a transmission electron microscope FEI Tecnai Biotwin 120kv.

For *in vivo i*EM, 6 months after transplantation, three rats were deeply anesthetized with pentobarbital and transcardially perfused with 0.1 M PBS followed by ice-cold 2% formaldehyde, containing 0.2% glutaraldehyde, in 0.1 M PBS, pH 7.4 for 1 hour. Brains were removed and washed in 0.1 M PBS. For *ex vivo* iEM the same fixation was performed for 30 min. Both, rat and human tissue were processed as described in **supplemental experimental procedures.**

### Electrophysiological recordings

For whole-cell patch-clamp recordings, 4 weeks-old human adult cortical organotypic slices were transferred to a recording chamber and were constantly perfused with carbogenated human artificial cerebrospinal fluid (hACSF, in mM: 129 NaCl, 21 NaHCO_3_, 10 glucose, 3 KCl, 1.25 NaH_2_PO_4_, 2 MgSO_4_ and 1.6 CaCl_2_, pH ~7.4) during the recordings. Recordings were performed with a HEKA EPC10 amplifier using PatchMaster software for data acquisition. The internal pipette solution contained (in mM): 122.5 K-gluconate, 12.5 KCl, 10 HEPES, 2.0 Na_2_ATP, 0.3 Na_2_-GTP, and 8.0 NaCl). Biocytin (1-3 mg/mL, Biotium) was dissolved in the pipette solution for *post-hoc* identification of recorded cells.

To study the ability to generate AP and its characteristics either a current ramp of 0-300 pA, or 10 pA current steps were applied at resting membrane potential (RMP) in the current clamp configuration. In voltage clamp mode, sodium and potassium currents were evoked by a series of 10 mV steps ranging from −70 mV to +40 mV. RMP was measured in current clamp mode immediately after establishing the whole-cell configuration. Input resistance (Ri) was calculated from a 10 mV pulse and monitored throughout the experiment. Data were analyzed offline with FitMaster and IgorPro software.

### Statistical analysis

Statistical analysis was performed using Prism 9 software (GraphPad). An unpaired t-test was used when data were normally distributed, whereas a Mann–Whitney U test was used when data did not pass the normality test. When different independent groups were compared, a one-way ANOVA plus Tukey’s multiple comparison tests were performed. Significance was set at P < 0.05. Data are mean ± SEM.

## ACKNOWLEDGMENTS

This work is supported by grants from the Swedish Research Council, Swedish Brain Foundation, Swedish Stroke Foundation, Region Skåne, Neurofonden, The Thorsten and Elsa Segerfalk Foundation, Rut och Erik Hardebo Foundation, and the Swedish Government Initiative for Strategic Research Areas (StemTherapy).

## AUTHOR CONTRIBUTIONS

S.P-T., O.L., and Z.K. conceived the project. R.M-C.; L.J.; C.A-M.; N.A.; O.T.; I.H.; E.M. and S.P-T. conducted experiments and analyzed data. S.P-T.; O.L.; and Z.K. wrote the manuscript. J.B. provided human material; R.M-C.; N.A.; E.M.; G.S. and O.T. were involved in the collection and/or assembly of data, data analysis, and interpretation. All authors reviewed and edited the manuscript.

