## Supplemental Information for "Oligodendrocytes in human iPS cell-derived cortical grafts remyelinate adult rat and human cortical neurons"

### SUPPLEMENTAL FIGURES AND TABLES

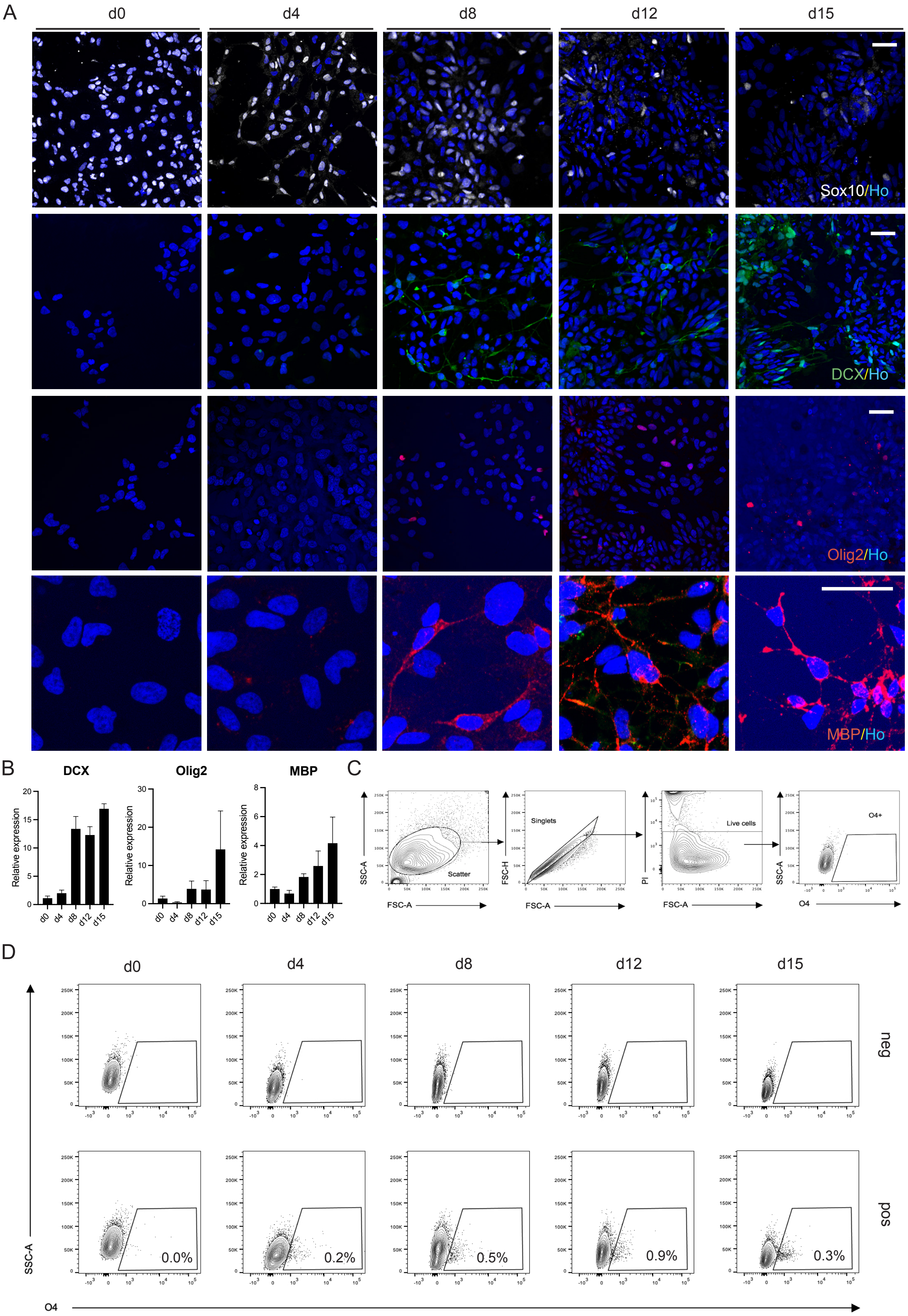


**Figure S1.** ***In vitro*** **differentiation of cortically-primed human lt-NES cells gives rise to a concomitant generation of neurons and oligodendrocytes.** (A) Confocal images of lt-NES cells at different time points of differentiation showing expression of the neuroectodermal and oligodendrocyte marker Sox10, the neuronal progenitor marker DCX, the pan-oligodendrocyte marker Olig2, and myelin basic protein MBP. (B) DCX, Olig2, and MBP changes in gene expression using RT-qPCR (n=3-4 replicates per time point). (C) Gating strategy used for flow cytometry analysis. (D) Flow cytometry analysis of live cells for $O4^{+}$ cells at 0, 4, 8, 12, and 15 days of differentiation. Negative control (without antibody) is shown on the top of the panel. Scale bar, 20 mm. Ho, Hoechst. Data are shown as mean ± SEM.

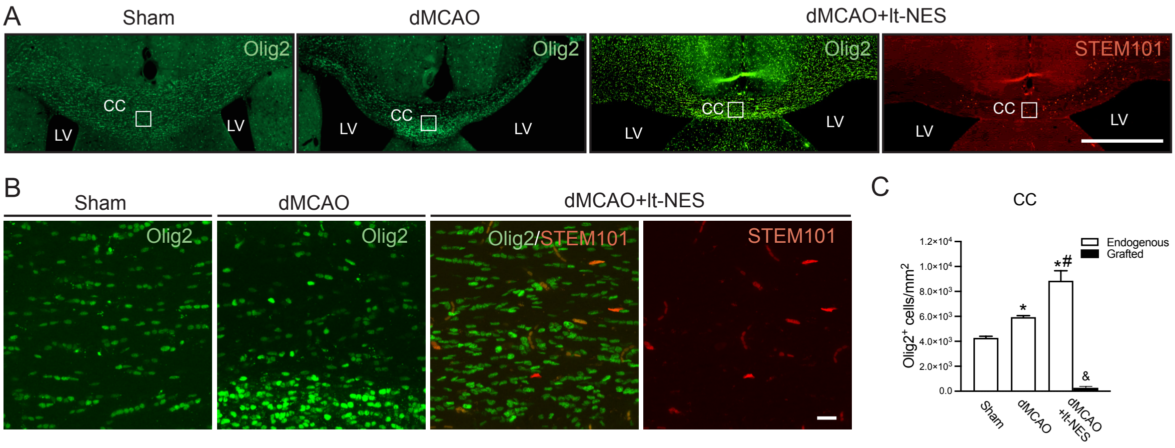


**Figure S2. Intracortical transplantation of human lt-NES cell-derived progenitors increases endogenous oligodendrogenesis in stroke-injured rat brains.** (A) Overview of Olig2$^{+}$ cells in the rat corpus callosum of sham-treated animals (Sham), and in animals subjected only to stroke (dMCAO) or stroke followed by transplantation (dMCAO+lt-NES). An overview of the location of graft-derived cells, using the human cell nuclear marker STEM101, is also included in the transplanted group. The white square depicts the area where images were taken and quantification was performed. Scale bar in A, 1 mm. CC: Corpus callosum. LV: Lateral ventricle. (B-C) Representative confocal images (B) and cell quantification (C) of Olig2$^{+}$ cells and STEM101$^{+}$ cells in the middle part of the corpus callosum. Scale bar in B, 20 µm. *p<0.05 vs. Sham. #p<0.05 vs. dMCAO. &p<0.05 vs. total number Olig2$^{+}$ in dMCAO+lt-NES cells. n=5 animals per group. Data are shown as mean ± SEM.

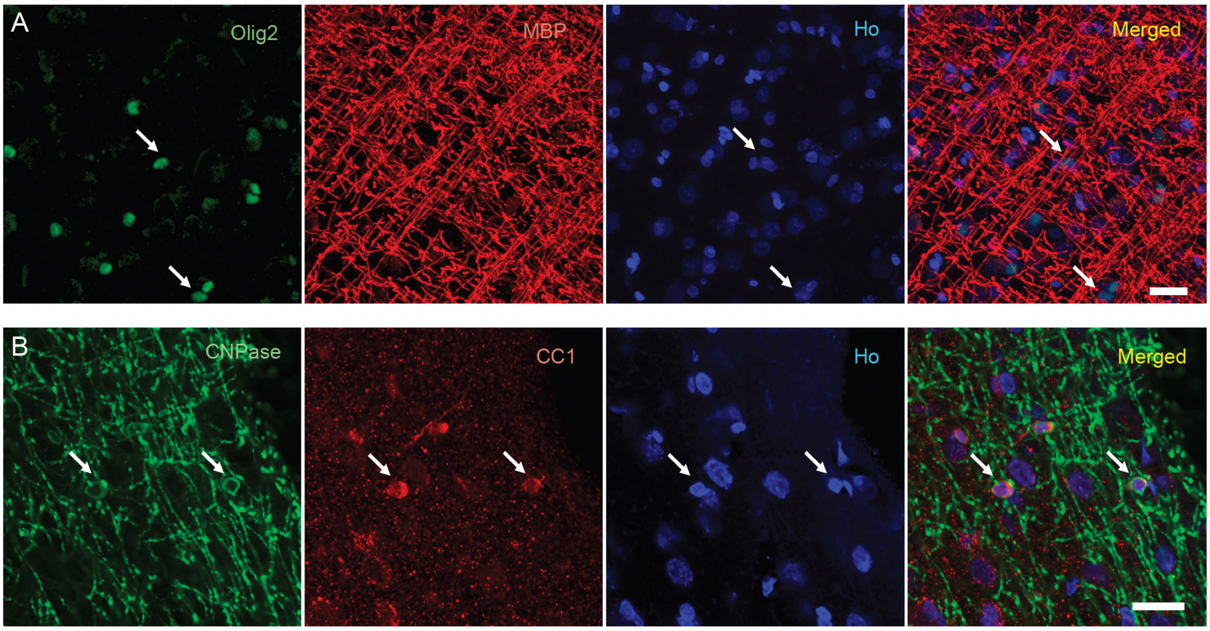


**Figure S3. Characterization of the oligodendrocyte population in acute human adult cortical tissue.** Representative confocal images of acute human cortical tissue showing the expression of different oligodendrocyte markers: (**A**) the pan-oligodendrocyte marker Olig2 and myelin basic protein (MBP), and (**B**) a marker for immature and mature oligodendrocytes (CNPase) and mature oligodendrocytes (CC1). Ho, Hoechst. Arrows indicate colocalization. Scale bar, 20 μm.

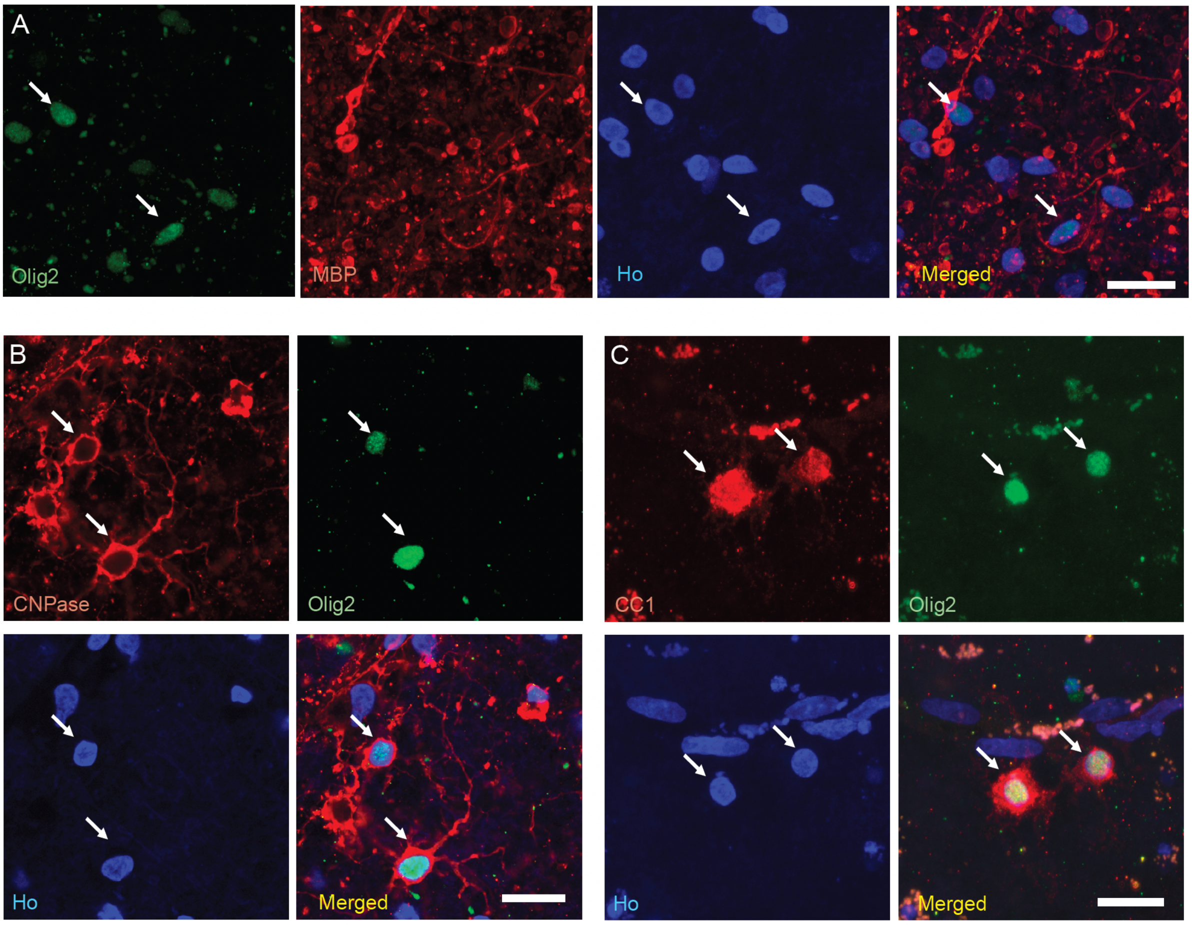


**Figure S4. The oligodendrocyte population is preserved in human adult cortical organotypic tissue after 4 weeks in culture.** Representative confocal images of 4 weeks-cultured human cortical tissue showing the expression of different oligodendrocyte markers: (**A**) the pan-oligodendrocyte marker Olig2 and myelin basic protein (MBP); (**B**) a marker for immature and mature oligodendrocytes (CNPase) and Olig2; and (**C**) a mature oligodendrocyte marker (CC1) and Olig2. Ho, Hoechst. Arrows indicate colocalization. Scale bar, 20 µm.

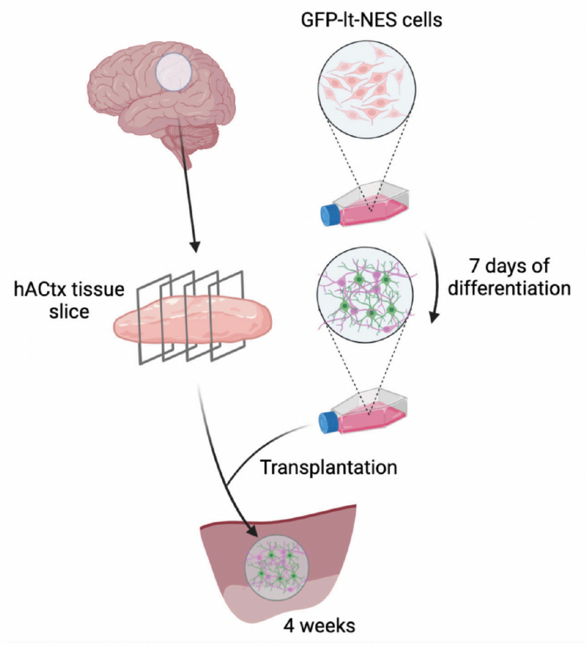


**Figure S5. Experimental design of lt-NES cell transplantation onto human adult cortical tissue.** Lt-NES cell-derived progenitors differentiated for 7 days are transplanted onto 300 μm slices of human adult cortical tissue cultured for a week. Coculture is maintained for 4 weeks before fixation. Generate using Biorender.

| **Antibody** | **Host species** | ***In vitro*** | ***In vivo*** | ***Ex vivo*** | **Notes** | **Company** |
|---|---|---|---|---|---|---|
| Primary Antibodies | | | | | | |
| CC1 | Mouse | 1:200 | 1:200 | 1:1000 | Antigen retrieval (AR) | Abcam |
| CNPase | Rabbit | - | 1:500 | 1:500 | AR | Abcam |
| DCX | Goat | 1:400 | - | - | | Santa Cruz |
| GFP | Chicken | - | - | 1:1000 | | Merk Millipore |
| MAP2 | Chicken | 1:500 | - | 1:1000 | | Abcam |
| MBP | Mouse | 1:1000 | 1:500 | 1:1000 | AR | Biolegend |
| NeuN | Rabbit | - | - | 1:1000 | | Abcam |
| Neurofilament | Chicken | 1:1000 | - | - | | Abcam |
| NG2 | Rabbit | - | 1:100 | - | | Merk Millipore |
| Olig2 | Rabbit | 1:500 | 1:500 | 1:500 | AR | Abcam |
| Sox10 | Goat | 1:200 | - | 1:500 | AR | Santa Cruz |
| STEM101 | Mouse | - | 1:500 | - | | Stem Cells |
| STEM121 | Mouse | - | 1:500 | - | | Stem Cells |
| Secondary Antibodies | | | | | | |
| 488 anti-Chicken | Donkey | 1:500 | - | 1:500 | | Jackson ImmunoResearch |
| 488 anti-Goat | Donkey | 1:500 | - | - | | Jackson ImmunoResearch |
| 488 anti-Mouse | Donkey | - | 1:500 | - | | Jackson ImmunoResearch |
| 488 anti-Rabbit | Donkey | 1:500 | 1:500 | 1:500 | | Jackson ImmunoResearch |
| Cy3 anti-Goat | Donkey | 1:500 | - | 1:500 | | Jackson ImmunoResearch |
| 647 anti-Chicken | Donkey | - | - | 1:500 | | Jackson ImmunoResearch |
| 647 anti-Mouse | Donkey | 1:500 | 1:500 | 1:500 | | Jackson ImmunoResearch |
| 647 anti-Rabbit | Donkey | 1:500 | 1:500 | 1:500 | | Jackson ImmunoResearch |

| | | | | | | |
|---|---|---|---|---|---|---|
| 647 Streptavidin | Donkey | - | - | 1:500 | | Jackson ImmunoResearch |

**Table S1.** List of primary and secondary antibodies.

### SUPPLEMENTAL EXPERIMENTAL PROCEDURES

#### Derivation of iPS cell-derived lt-NES cells

Human dermal fibroblasts from healthy adult donors were subjected to Sendai virus transduction with the reprogramming factors Oct4, Sox2, KLF4, and c-MYC (CytoTune iPS 2.0 Sendai Reprogramming kit, Invitrogen). Colonies were picked to establish iPS cell lines using mTeSR medium (Invitrogen). For neural induction, iPS cells were split and colonies were gently resuspended in embryoid body (EB) medium (Dulbecco's modified Eagle medium/F12 [DMEM/F12], 10% KSR, 2-Mercaptoethanol [1:1000], nonessential amino acids [NMEAA] [1:100], Glutamine [1:100]) with Rock Inhibitor (1:1000), 3 μM Dorsomorphin (Sigma-Aldrich) and 10 μM SB431542 (Sigma-Aldrich). On day 5, EBs were collected and plated on poly-L-ornithine/ laminin-coated plates in an EB medium with 3 μM Dorsomorphin and 10 μM SB431542. On day 6, the media was changed to N2 medium (DMEM-F12 [without Hepes + Glutamine], N2 [1:100], Glucose [1.6 g/L]) supplemented with 1 μM Dorsomorphin and 10 ng/mL bFGF. Neural rosettes were carefully picked, six days later, and grown in suspension in an N2 medium with 20 ng/mL bFGF. On day 14, neural rosette spheroids were collected, and the small clumps obtained were grown in adhesion on poly-L-ornithine/laminin-coated dishes in the presence of 10 ng/mL bFGF, 10 ng/mL EGF (both from Peprotech) and B27 (1:1000, Invitrogen). The lt-NES cell line was routinely cultured and expanded on 0.1 mg/mL poly-L-ornithine and 10 mg/mL laminin (both from Sigma)-coated plates into the same media supplemented with FGF, EGF, and B27 and passaged at a ratio of 1:2 to 1:3 every second to the third day using trypsin (Sigma).

#### Distal middle cerebral artery occlusion and cell transplantation

Animals were housed in individually ventilated cages under standard temperature and humidity conditions and a 12-h light/dark cycle with free access to food and water.

The focal ischemic injury was induced in the somatosensory cortex by distal middle cerebral artery occlusion (dMCAO) as described previously (Chen et al., 1986; Oki et al., 2012). Briefly, Animals were anesthetized with isoflurane (3.0% induction; 1.5% maintenance) mixed with air and the temporal bone was exposed. A craniotomy of 3 mm was made, the dura mater was carefully opened, and the cortical branch of the middle cerebral artery was ligated permanently by suture. Both common carotid arteries were isolated and ligated for 30 min. After releasing common carotid arteries, surgical wounds were closed.

Intracortical transplantation of cortically fated $GFP^+$ lt-NES cell-derived progenitors was performed stereotaxically 48 h after dMCAO as described previously (Palma-Tortosa et al., 2020; Tornero et al., 2013). Briefly, on the day of surgery, cortically primed cells on their third day of differentiation were resuspended to a final concentration of 1 x $10^5$ cells/μL in cytocon buffer. A volume of 1 μL was injected at two sites with the following coordinates: anterior/posterior: +1.5 mm; medial/lateral: 1.5 mm; dorsal/ventral: - 2.0 mm; and anterior/posterior: + 0.5 mm; medial/lateral: 1.5 mm; dorsal/ventral: -2,5 mm.

#### Organotypic cultures of the adult human cortex

The surgically resected tissue was instantly kept in ice-cold modified human artificial cerebrospinal fluid and sliced on a vibratome (Leica VT200S). Slices of 300 μm thickness were kept in a rinsing medium until they were fixed with 4% formaldehyde for acute characterization of the tissue or transferred to cell culture inserts containing Alvetex scaffold membranes (Reinnervate) in 6 well plates filled with human adult cortical media (Brain Phys medium [without phenol red] supplemented with B27 [1:50], glutamax [1:200] and gentamycin [50 mg/mL]), and incubated in 5% CO2 at +37ºC. The organotypic slices were maintained in culture for 1 week before transplantation of $GFP^+$ lt-NES cells and kept 4 weeks more before characterization of grafted cells. Non-transplanted slices maintained for 4 weeks in culture were used for characterization by immunohistochemistry and electrophysiological recordings of the neural populations in cultured tissue.

**Immunocytochemistry and Immunohistochemistry**

Cortically fated lt-NES cells plated on glass coverslips were fixed at different time points of differentiation in 4% formaldehyde (Sigma) for 20 min at room temperature. Cells were permeabilized with 0.025% Triton X-100 in 0.1 M potassium phosphate buffered saline (KPBS) and blocked with 5% of normal donkey serum (NDS) for 45 min at room temperature. Afterward, primary antibodies (**Table S1**) diluted in blocking solution were applied overnight at +4 ºC followed by 3 rinses with KPBS. Fluorophore-conjugated secondary antibodies (**Table S1**) (1:500, Jackson Laboratories) diluted in blocking solution were applied for 2 h at room temperature. Subsequently, cells were rinsed 3 times and nuclei were stained with Hoechst (Molecular Probes or Jackson Laboratories) for 10 min at room temperature. Stained glass coverslips were mounted on slides with Dabco (Sigma) mounting media.

Regarding immunohistochemistry in rat slices, the stored sections were rinsed 3 times with KPBS and incubated in a blocking solution for 1 h (10% NDS and 0.25 Triton X-100 in 0.1 M KPBS [TKPBS]). The rest of the procedure follows that outlined above for cell cultures. The list of primary and secondary antibodies used can be found in **Table S1.**

For staining in human organotypic cultures, slices were fixed with 4% formaldehyde overnight at + 4 ºC and rinsed 3 times with KPBS for 15 min. Slices were then incubated overnight at + 4 ºC in permeabilization solution (0.02% Bovine serum Albumin [BSA], 1% Triton X in PBS), and overnight at +4 ºC in blocking solution (KPBS, 0.2% Triton X-100, 1% BSA, Sodium azide [NaN3] [1:10000] and 10% NDS). Primary antibodies were diluted in a blocking solution and incubated for 48 h at +4 ºC. Secondary antibodies were then diluted in a blocking solution and applied for 48 h at +4 ºC. Slices were washed 3 times with KPBS and incubated in Hoechst for 2 h at room temperature. Finally, mounted with Dabco after rinsing with deionized water.

Some stainings required antigen retrieval (**Table S1**) before the permeabilization step. Cells, rat sections, or human organotypic cultures were incubated with sodium citrate pH 6.0 Tween 0.05%, for 30 min (for cells and sections) or 2 h (for organotypic cultures) at +65 ºC.

**Immuno-electron microscopy**

For iEM in rat tissue, frontal 100 µm sections of the whole brain were cut on a Vibratome VT1000A (Leica, Germany). The sections were cryoprotected, freeze-thawed in liquid nitrogen, and incubated overnight in primary goat anti-GFP antibody (1:500, Novus Biologicals) at +4 ºC. The tissue was then incubated at room temperature for 2 h with biotinylated rabbit anti-goat secondary antibody (1:200, DakoCytomation), and avidin-biotin-peroxidase complex (ABC) (Vector Laboratories) followed by 3,3′-diaminobenzidine tetrachloride (DAB) and 0.015% hydrogen peroxide. Following the DAB reaction, sections were processed for iEM. DAB-immunostained sections of rat brain tissue and human organotypic cultures were postfixed in 1% osmium tetroxide in 0.1 M PBS, dehydrated in a graded series of ethanol and propylene oxide, and flat-embedded in Epon. Ultrathin sections were cut with a diamond knife. For post-embedding immunogold labeling of GFP, ultrathin sections were incubated overnight in primary goat anti-GFP antibody (1:500, Novus Biologicals) at +4 ºC. A secondary antibody (donkey anti-rabbit IgG conjugated to 12 nm colloidal gold; Jackson Laboratories) diluted 1:20 in 0.1% BSA in PBS was added for 1.5 h, then washed with PBS. Sections were then fixed with 2% glutaraldehyde, then washed with PBS followed by $dH_2O$. Sections were stained with uranyl acetate and lead citrate. Ultrathin sections were examined and photographed using a transmission electron microscope JEM- 100CX (JEOL, Japan).
